# Limbal Niche Cells in Three-Dimensional Matrigel Induced Dedifferentiation of Mature Corneal Epithelial Cells toward a Progenitor State

**DOI:** 10.1101/2020.06.01.128629

**Authors:** Hui Zhu, Wei Wang, Lingjuan Xu, Menglin Jiang, Yongyao Tan, Yunming Wang, Yaa-Jyuhn James Meir, Guigang Li

## Abstract

**Purpose:** To investigate the possibility and the key factors of stably committed mature corneal epithelial cells dedifferentiate into corneal epithelial stem cells *in vitro*.

**Methods:** Mature cornea epithelia cell (MCEC) sheets or limbal epithelial progenitor cell (LEPC) sheets were isolated from central corneas or limbal segments by Dispase II and further digested with 0.25% trypsin/1 mM EDTA (T/E) to yield single cells. Limbal niche cells (LNC) were isolated from the limbal stroma by collagenase A and expanded on 5% Matrigel coated plastic. Single MCECs were seeded on 50% Matrigel with or without LNC culturing for 10 days, regarding as three-dimensional MCEC (3D-MCEC) group or three-dimensional MCEC+LNC (3D-MCEC+LNC) group. Expression of CK12, p63α, PCK, Vimentin were analyzed with immunofluorescence staining.

**Results:** The expression of mature cornea epithelial marker (CK12) in MCEC was higher than that in LEPC (P=0.020) but epithelial stem cell marker (p63α) was lower than that in LEPC (P=0.000). When seeded in 3D Matrigel, single MCEC cells could form spheres within 72 hours, and the expression of CK12 reduced (P=0.005) and the expression of p63α also reduced to zero (P=0.000) compared to MCEC. Serial passages of LNC which were expanded in coated Matrigel could form spheres in 3D Matrigel. After mixing MCECs with LNC, rounder spheres emerged within 24 hours which consisted of both epithelia cells (PCK^+^/Vim^-^) and LNC (PCK^-^/Vim^+^). Moreover, epithelia cells in 3D-MCEC+LNC group expressed less CK12 and more p63α than those in MCEC group (P=0.043, 0.000). Besides, the diameter of spheres in 3D-MCEC+LNC group were larger than that in 3D-MCEC group (P=0.000).

**Conclusion:** Human LNC and three-dimensional Matrigel could induce the dedifferentiation of mature corneal epithelial cells into corneal epithelial stem cells.

## INTRODUCTION

Corneal disease is the second major cause of blindness worldwide, which attributes to 3.2% of global blindness^1^. In China, there are approximately 3 million blindness due to corneal diseases, with 200,000 cases increasing every year^2^. Limbal stem cell deficiency (LSCD) is one of the most common debilitating eye disorders to cause corneal blindness. As we all know, limbus of the cornea forms a border between the corneal and conjunctival epithelium and its limbal stem cells (LSCs), located in limbal palisades of Vogt, are known as stem cells to generate mature corneal epithelial cells (MCEC), maintaining and repairing the intact cornea epithelial layer^3^. The quiescent state, self-renewal, proliferation and terminal differentiation of LSC are all regulated by limbal microenvironment^4^. However, LSCs could be destroyed and lost under various etiologies, which can result in a delay of corneal epithelial regeneration, overgrowth of conjunctival-derived epithelial cells, corneal stromal neovascularization, and corneal opacity^5, 6^. Studies have confirmed that patients with one-eye LSCD can be cured by autologous limbal stem cell transplantation^7^, but patients with binocular LSCD who have completely lost their autologous LSC can only be adopted allograft LSC transplantation or autologous cultivated replacement cell transplantation at present^8^. The main defect of allogenic LSC transplantation is severe rejection, while autologous replacement cells, such as oral mucosal epithelial cells^9, 10^, acquire some of the corneal epithelial-like characters, they have higher angiogenic potential and the uncertainty of long-term transplantation effects.

Regeneration of injured tissues is usually associated with cell fate plasticity, in which cells deviate from their normal lineage paths. Therefore, it is becoming gradually evident that this plasticity can provide strategies to restore damaged stem cells^11^. Pioneering work that began in the late 1800s had found that newts’ retinas, lenses and even limbs could regenerate, suggesting that some vertebrates retain outstanding cell fate plasticity. In adult newts, mature cells can dedifferentiate and become proliferative while maintaining their original lineage commitment and forming cells loyal to the fate of the original progenitor cell^12^. In recent years, studies using lineage tracing technology have gradually proved the phenomenon of original cell fate changes caused by injury in many mammalian tissues, among which the most important discovery is the reverse differentiation of epithelial cells under pathological conditions^11^. In 2013, Purushothama Rao Tata et al.^13^ demonstrated that highly differentiated airway epithelial cells can revert into stable and functional stem cells in vivo. After ablation of airway epithelial stem cells, it demonstrated that the luminal secretory cells had dedifferentiated into basal stem cells using lineage tracing, which not only show indistinguishable morphology from stem cells and express stem cell markers (CK5, p63), but also could proliferate and differentiate to produce new epithelial cells and repair damaged epithelial areas. Moreover, single secretory cells could dedifferentiate into multipotent stem cells when they were cultured in 3D Matrigel environment in vitro. In 2018, Waseem Nasser et al.^14^ discovered that following deletion of limbal stem cell boundary, mature corneal epithelial cells could dedifferentiate into K15-GFP^+^ stem cells and reform the tissue boundary in the presence of an intact niche. However, niche destruction abolished K15-GFP^+^ recovery and induced severe LSCD, suggesting that dedifferentiation of corneal cells into K15-GFP^+^ LSCs requires an intact limbal stroma. This phenomenon of cell fate plasticity provides new therapeutic strategies for repairing damaged or lost stem cells, and the success of dedifferentiation is dependent on the original stem cell microenvironment. Therefore, our study investigated whether MCEC can be reversely differentiated into LSC in a simulated environment in vitro.

The components of LSC microenvironment^15^ include extracellular Matrigel, pigment, stromal cells, blood vessels and nerves, etc. In previous studies, we isolated limbal niche cells (LNC) from the limbus niche of cornea, which express Vimentin and other embryonic stem cell markers, such as Oct4, Sox2, Nanog but don’t express PCK, and it is a group of endothelial cells progenitors^16^ and mesenchymal Stem cells (MSC)^17^ progenitors. We also concluded that LNC were a more powerful resource than bone marrow mesenchymal stem cell (BMMSC) to prevent LSCD in an alkali burn subconjunctival rabbit model partially due to increased activation of SCF signaling^18^. It is suggested that LNC are indispensable supporting cells for LSC microenvironment.

The three-dimensional (3D) Matrigel environment can induce the reverse differentiation of mature cells into stem cells^19^. In previous study, serial passages on 5% coated Matrigel resulted in rapid expansion of LNC, of which the expression of embryonic stem cell markers (Oct4, Sox2, Nanog, Rex1, SSEA4) and vascular progenitor cell markers (flk-1, CD34, CD31) reduced gradually but could be restored after reseeding in 3D Matrigel in MESCM^20^. Moreover, 3D Matrigel could induce LNC differentiated into vascular endothelial cells and pericyte cells stabilizing the tube network formed by HUVEC^16^. After Limbal epithelial progenitor cell (LEPC) co-cultured with LNC in 3D Matrigel for 10 days, resultant epithelial spheres mixed with spindle cells expanded in MESCM could express more epithelial stem cell marker (p63α) and less corneal specific differentiation mature marker (CK12)^20^. In our present research, we found that MCEC have the possibility to dedifferentiate into corneal epithelial stem cell (CESC) with the help of LNC in 3D Matrigel, which could be a potential novel therapy of LSCD.

## MATERIALS AND METHODS

### Tissue Isolation and Cell Culturing

Donor corneas from 18 to 60 years old were obtained from the Red Cross Eye Bank of Wuhan City, Tongji Hospital (Hu Bei, China) and managed in accordance with Declaration of Helsinki and approved by Tongji Ethics Committee. The central or limbal explants were digested with 10 mg/ml Dispase II at 4 °C for 16 hours to generate intact central and limbal epithelial sheets. Then the epithelial sheets were digested further with 0.25% trypsin and 1mM EDTA (T/E) at 37°C for 15 minutes to yield single epithelial cells.

Human limbal niche cells (LNC) were isolated and cultured as previously prescribed ^20^. Limbal explants were digested with collagenase A at 37°C for 8-10 hours to generate clusters containing the entire limbal epithelial sheet with subjacent stromal cells. Then the clusters were digested further with T/E at 37°C for 15 minutes to obtain single cells before being seeded at the density of 1×10^4^ per cm^2^ in 6-well plates either on coated Matrigel in ESCM containing 10 ng/ml LIF and 4 ng/ml bFGF (MESCM). ESCM is made of knockout DMEM supplemented with 10% knockout serum, 5 μg/ml insulin, 5 μg/ml transferrin, 5 ng/ml selenium, 50 μg/ml gentamicin, and 1.25 μg/ml amphotericin B. Upon 80-90% confluence, they were passaged serially at the density of 5× 10^3^ per cm^2^. All materials used for cell isolation and culturing are listed in Supplementary Table S1.

### Culturing in Three-Dimensional Matrigel

As reported previously^17^, three-dimensional (3D) Matrigel was prepared by adding 150 μl of 50% Matrigel (diluted in MESCM) per chamber of an 8-well chamber slide following incubation at 37°C for 30 minutes. Single central epithelial cells obtained from Dispase II-isolated epithelial sheets were seeded at the total density of 12×10^4^ /cm^2^ in 3D Matrigel. Serial passage 4 LNC which were expanded in coated Matrigel were harvested and then seeded in 3D Matrigel at the total density of 12×10^4^ /cm^2^. And single central epithelial cells were mixed at a ratio of 4:1 with the P4 LNC and seeded at the total density of 12×10^4^ /cm^2^ in 3D Matrigel. After 10 days of culture in MESCM, the resultant sphere growth was collected by digestion of Matrigel with 10 mg/ml Dispase II at 37°C for 2 hours. Then the spheres were digested further with T/E at 37°C for 15 minutes to yield single cells.

### Immunofluorescence Staining

Central or limbal epithelial sheets obtained by Dispase II digestion were cryosectioned to 6 μm. Single cells and spheres were prepared for cytospin using Cytofuge at 1000 rpm for 8 minutes (StatSpin, Inc., Norwood, MA). For immunofluorescence staining, the samples were fixed in 4% paraformaldehyde, and then permeated with 0.2% Triton X-100 in PBS for 15 minutes, and blocked with 2% BSA in PBS for 1 hour before being incubated with primary antibodies overnight at 4°C. After washing with PBS, corresponding secondary antibodies were incubated for 1 hour using appropriate isotype-matched nonspecific IgG antibodies as controls. The nucleus was counterstained with DAPI before being analyzed with a Zeiss LSM 700 confocal microscope (LSM700; Carl Zeiss). The quantitation was done by ImageJ. Detailed information about primary and secondary antibodies and agents used for immunostaining is listed in Supplementary Table S2.

### Statistical Analysis

The experimental data were represented as means ± SD and compared using the appropriate version of student’s *t*-test. Test results were reported as two-tailed *P* values, where *P* < 0.05 was considered statistically significant. Statistical analysis was performed using Graphpad Prism 8.0 and ImageJ.

## RESULTS

### Epithelial cells from the central cornea express more CK12while less p63α than that from limbus

According to the previous methods^16, 17, 20^, we used limbal corneal epithelium progenitor cell (LEPC) isolated by Dispase II as control group. Epithelial cells were isolated with Dispase II from central cornea and limbus, either were prepared for frozen cross section (A, scale bar=20μm) or further digested with T/E to get single cells before fixed on slides with cytospin (B), and then both were labeled with CK12 and p63α immunofluorescence staining separately. The positive rate were calculated and compared between groups based on single cell counting from B (C). 100% of the epithelial cells from the central cornea expressed CK12, while 36.96±3.96% of them expressed p63α, which mainly localized in the basal layer of the cell sheet. In contrast, 26.43±18.10% of the epithelial cells from the limbus expressed CK12, which localize mainly at the outer layer of the cell sheet, while84.73±4.67% of the cells expressed p63α, which localized mainly at the basal layer of the cell sheet. The CK12 positive rate of epithelial cells from the central cornea was statistically higher than that from the limbus (P=0.020), while p63 positive rate of epithelial cells from the central cornea was statistically lower than that from the limbus (P=0.000) (C)

### MCEC could form spheres in 3D Matrigel, with reduced expression of CK12 and p63α

Previous research has shown that LEPC digested by Dispase II culturing in 3D Matrigel alone, when choosing MESCM as culture medium, could form spheres consisting of the PCK+ epithelial cells ^17^. Therefore, we put mature central epithelial cell (MCEC) digested by Dispase II from central cornea culture in 3D Matrigel alone (3D-MCEC group). It showed that MCECs could also gradually gather within 72 hours into the spherical structure. However, the cells were spread unevenly, and spheres were in irregular shape on the day10. (Fig.2 A) Immunofluorescence double staining was performed on all the unaggregated cells and aggregated cell spheres in 3D Matrigel on day10. The positive expression rate of CK12 was 24.06±8.85%, but p63α expression was negative in both the spheres and the unaggregated cells. (Fig.2 B) Comparing with MCEC group, the positive expression rate of CK12 decreased in 3D-MCEC group (P=0.005), which suggested that 3D Matrigel can reduce the maturity of MCEC. (Fig.2 C) However, the positive expression rate of p63α also decreased in 3D-MCEC group (P=0.000)

**Figure 1.**
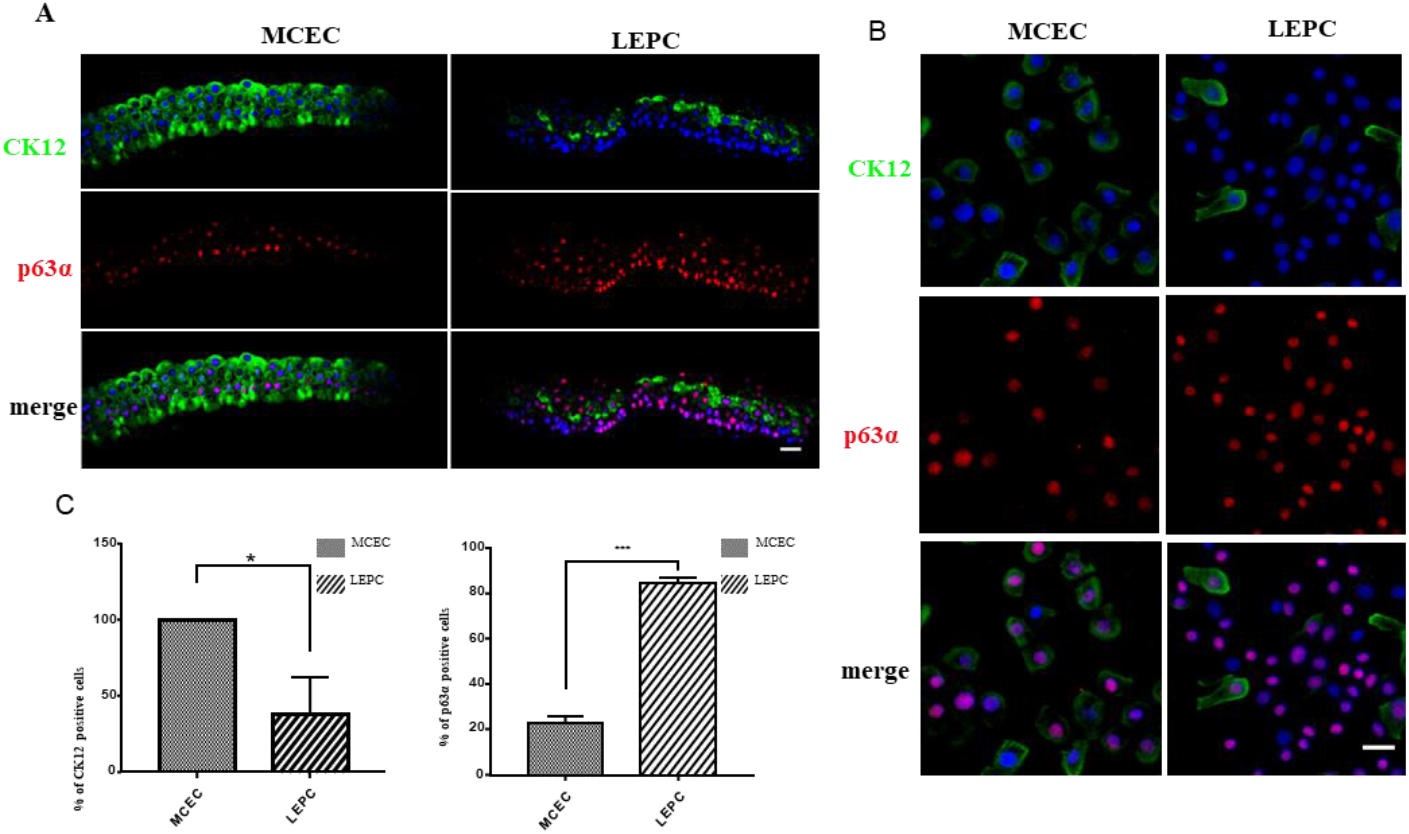
Epithelial cells from the central cornea express more CK12 while less p63α than that from limbus. Epithelial cells were isolated with Dispase II from central cornea and limbus, either were prepared for frozen cross section (A, scale bar=20μm) or further digested with T/E to get single cells before fixed on slides with cytospin (B), and then both were labeled with CK12 and p63α immunofluorescence staining separately. The positive rate was calculated and compared between groups based on single cell counting from B (C). 100% of the epithelial cells from the central cornea expressed CK12, while 36.96±3.96% of them expressed p63α, which mainly localized in the basal layer of the cell sheet. In contrast, 26.43±18.10% of the epithelial cells from the limbus expressed CK12, which localize mainly at the outer layer of the cell sheet, while84.73±4.67% of the cells expressed p63α, which localized mainly at the basal layer of the cell sheet. The CK12 positive rate of epithelial cells from the central cornea was statistically higher than that from the limbus (P=0.020), while p63 positive rate of epithelial cells from the central cornea was statistically lower than that from the limbus (P=0.000) (C)

**Figure 2.**
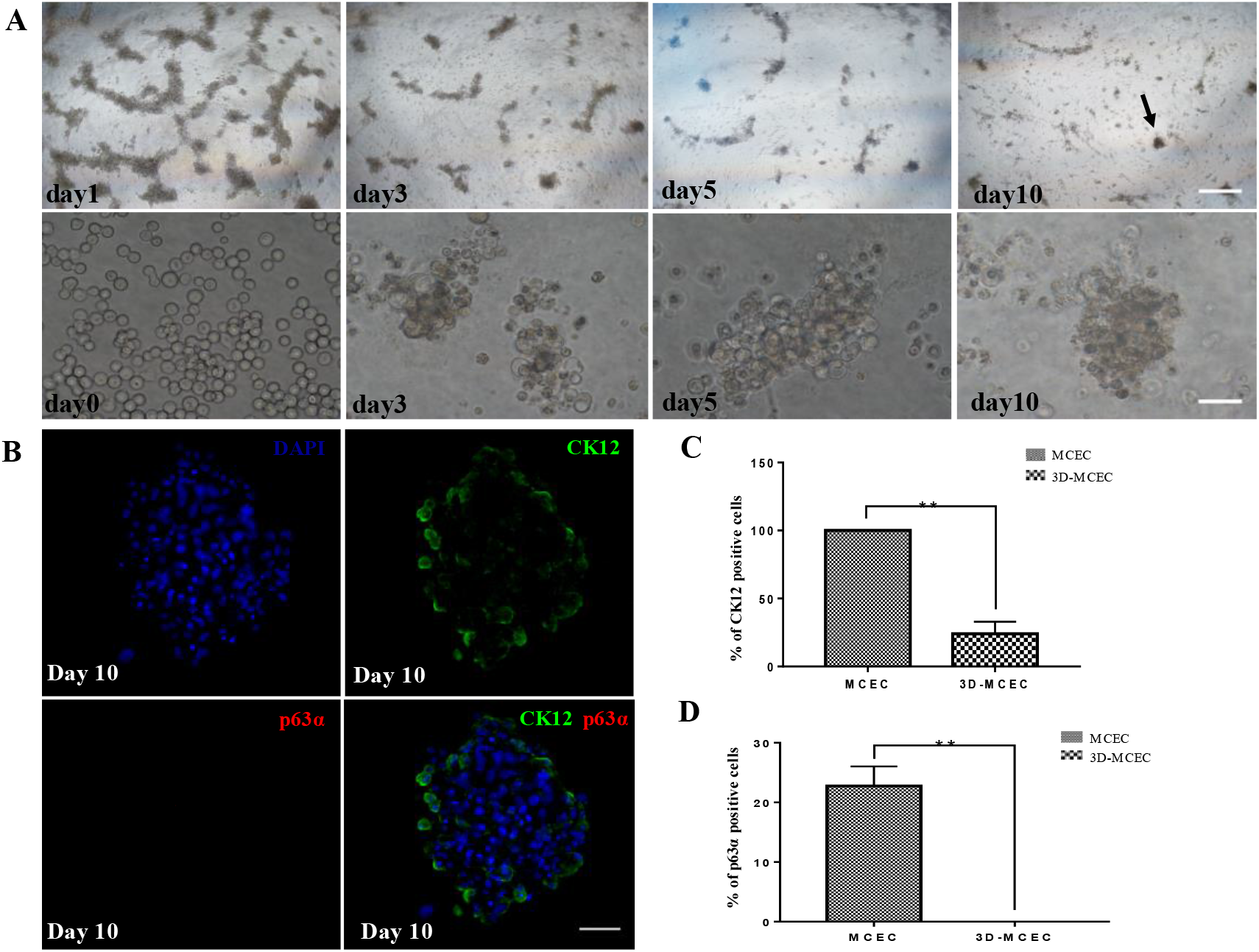
MCEC cultured in 3D Matrigel alone and the expression changes of CK12 and p63. MCEC sheets were digested by Dispase II then followed by T/E method to obtain single cells which were seeded in 50% 3D Matrigel at a density of 12×10^4^/cm^2^ culturing as 3D-MCEC group. Single cells in 3D-MCEC group could accumulate gradually within 72 hours but the cells were not so evenly distributed that only a few cells gathered into spheres on day10. (Fig.2A, black arrow points to sphere, upper row scale bar=500μm, lower row scale bar=50μm). Immunofluorescence double staining showed 24.06±8.85% of the cells in the 3D-MCEC group sphere expressed CK12 on day 10, but no cells expressed p63 (Fig.2B, scale bar=50μm). The expression CK12 of spheres in 3D-MCEC group was lower than that of single cells in MCEC (P=0.005) but expression of p63α also was lower than that in MCEC (P=0.000). (Fig.2C D)

### LNC could expand in coated Matrigel and form spheres in 3D Matrigel

LNC were isolated and expanded from limbal cluster by Collagenase A digestion followed with T/E and cultured in different environment. Immunofluorescence staining on single cells from limbal clusters show that there were mixed cells with expression of CK12+ epithelial cells, p63α+ limbal progenitor cells and CK12-/p63α-LNC.(Fig.3A) Comparing the two different microenvironment culturing LNC, it showed that single cells seeded in 5% Matrigel could isolate and expand spindle cells, which was demonstrated as LNC in previous study^20^. (Fig.3B) Interestingly, it showed that P4 LNC seeded in 50% 3D Matrigel could aggregate within 24h and form huge spheres on day10, suggesting that 3D environment could have an influence on LNC. (Fig.3C)

**Figure 3.**
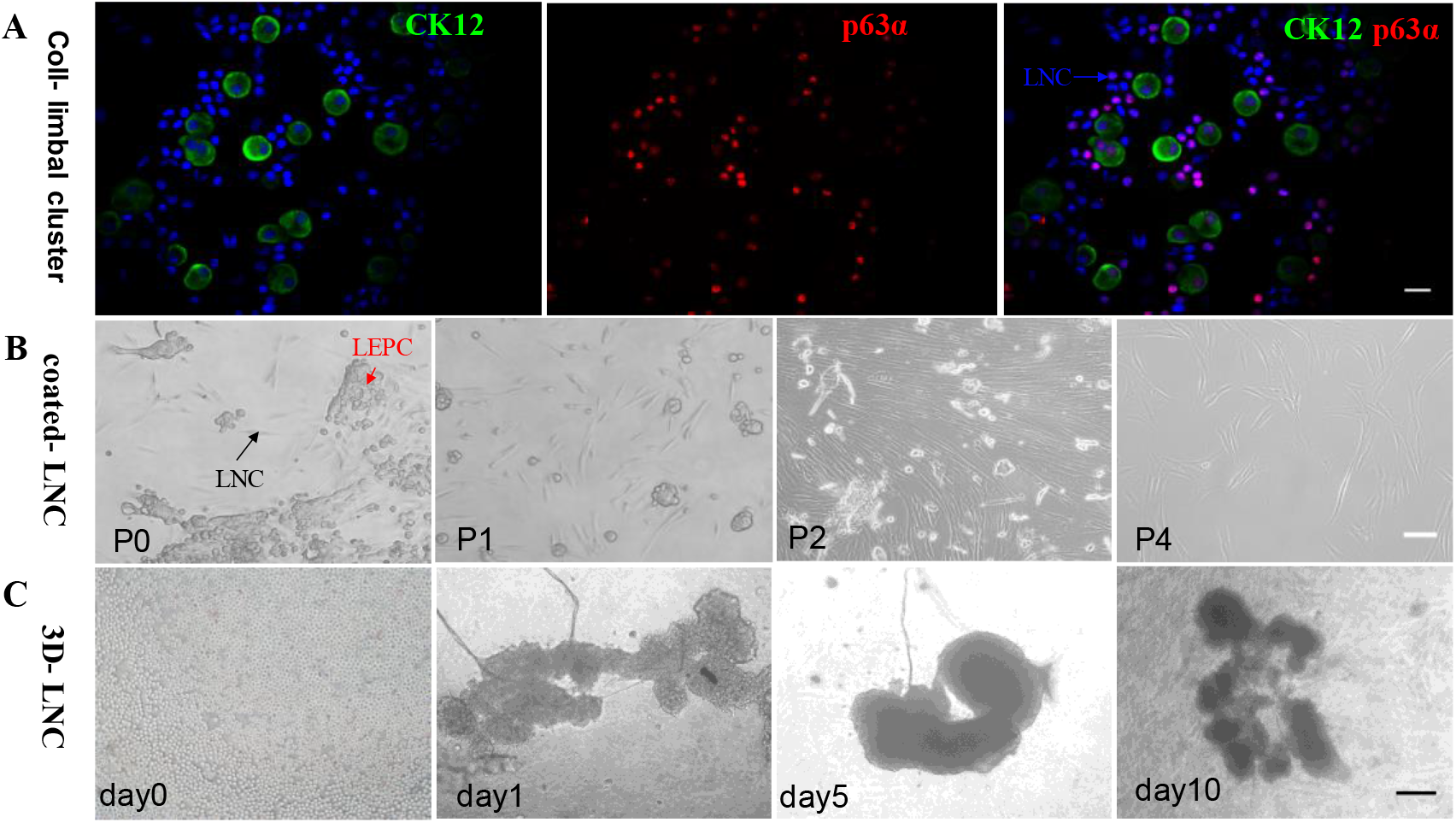
Expansion of LNC cultured in coated and aggregation of P4LNC cultured in3D Matrigel. Collagenase A-isolated limbal clusters followed by T/E could generate single cells. Immunofluorescence staining on these single cells showed that there were mixed cells with expression of CK12+ epithelial cells, p63α+ LEPCs, and PCK-/p63α-LNC, consistent with our previous study^20^. (Fig.3A, blue arrow points to LNC, scale bar A=20μm) Single cells derived from Collagenase A-isolated limbal clusters followed by T/E were seeded on 5% coated Matrigel at density of 1×10^5^/cm^2^ in MESCM. Spindle cells, demonstrated as LNC in our previous study^17, 20^, emerged in P0, distinguished from LEPCs, and then proliferated and became dominant after P2. (Fig.3B, red arrow points to LEPCs, black arrow points to LNC, scale bar A=100μm) P4 LNC were seeded in 50% 3D Matrigel could aggregate and form huge spheres. (Fig.3C, scale bar C=200μm)

### MCEC mixing with LNC could be united in spheres growth in 3D Matrigel

Previous studies^17^ have shown that limbal clusters isolated by collagenase A method contained 20% of the PCK-/Vim+ limbal niche cells (LNC). Therefore, we mixed single cells of MCEC with LNC at a ratio of 4:1 and cultured them on 3D Matrigel (3D-MCEC+LNC group). It showed that all the cells began to assemble into spherical structures within 24 hours. On the day10 of culturing, the cell spheres were evenly distributed. (Fig.4 A) Then all spheres of 3D-MCEC+LNC group were collected for immunofluorescence staining. PCK+/Vim- and PCK-/Vim+ cells could be seen in the cell spheres, suggesting that the spherical shape was composed of epithelial and stromal cells, which demonstrated that MCEC and LNC could be reunited in 3D Matrigel environment. (Fig.4 B)

**Figure 4.**
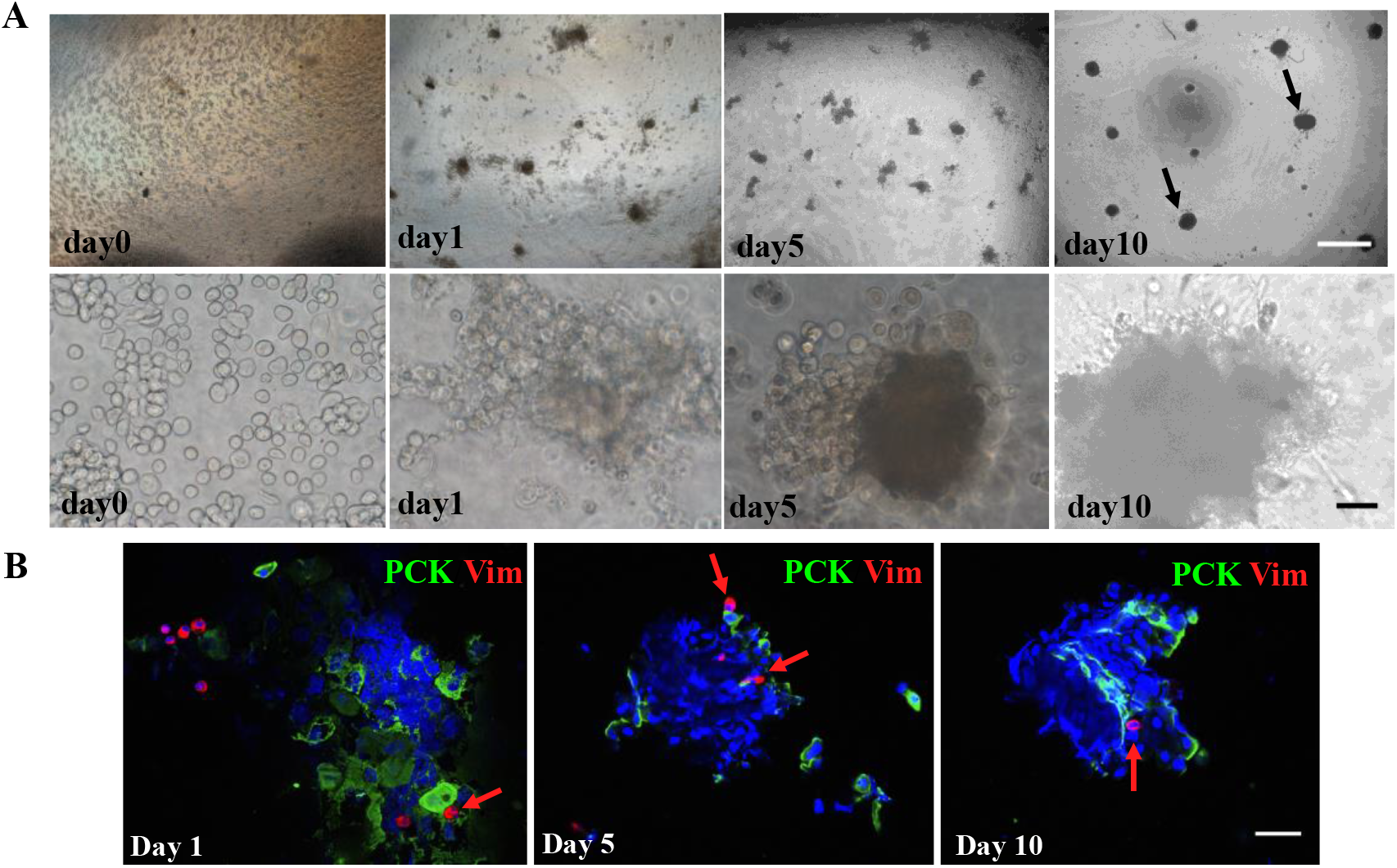
MCEC mixed with LNC could form more spheres in 3D Matrigel, with epithelial cell and LNC united. Single mature central epithelial cells were obtained from Dispase II -isolated MCEC sheets followed by T/E, and mixed with P4 LNC at a ratio of 4:1 and then seeded in 50% 3D Matrigel at a density of 12×10^4^/cm^2^ culturing as 3D-MCEC+LNC group. All Cells in 3D-MCEC+LNC group could assemble into spheres within 24 hours, with more spheres evenly distributed on day10. (Fig.4A, black arrow points to sphere, upper row scale bar=500μm, lower row scale bar=50μm). All spheres of 3D-MCEC+LNC group were collected using Dispase II method for immunofluorescence staining on the day1, day5 and day10. Immunofluorescence double staining of spheres showed that epithelial cells (PCK+/Vim-) and LNC (PCK-/Vim+) could be seen gradually gathered in spheres. (Fig.4B, red arrow points to LNC, scale bar=50μm)

### 3D-MCEC+LNC group expressed less CK12 while more p63α than that in 3D-MCEC group

On the day10 of culturing, all spheres in 3D-MCEC+LNC group were collected by Dispase II digestion. Immunofluorescence staining of cell spheres showed that there were some cells expressed p63α and other cells expressed CK12 concluded in one sphere.(Fig.5A upper row) Then we digested the spheres into single cells with T/E. The immunofluorescence staining of single cells from the sphere showed that the positive rate of CK12 and p63α was 52.95±20.99% and 35.58±8.11% respectively.(Fig.5A lower row) Comparing with the MCEC group, the positive rate of CK12 in the 3D-MCEC+LNC group reduced (P=0.000) and p63α expression increased (P=0.043), among which the proportion of p63α+/CK12-cells were 11.51±3.76%.(Fig.5B) we also compared three groups (MCEC, 3D-MCEC, 3D-MCEC+LNC). However, there was no statistically significant difference in CK12 positive rate between the 3D-MCEC group and the 3D-MCEC+LNC group (P=0.051), indicating that the 3D Matrigel environment had the same effect on CK12 expression reduction in the two groups. (Fig. 6 A) Even though there was no p63α expression in 3D-MCEC group, 3D-MCEC+LNC expressed more p63α than MCEC group (P=0.043) (Fig. 6 B)

**Figure 5.**
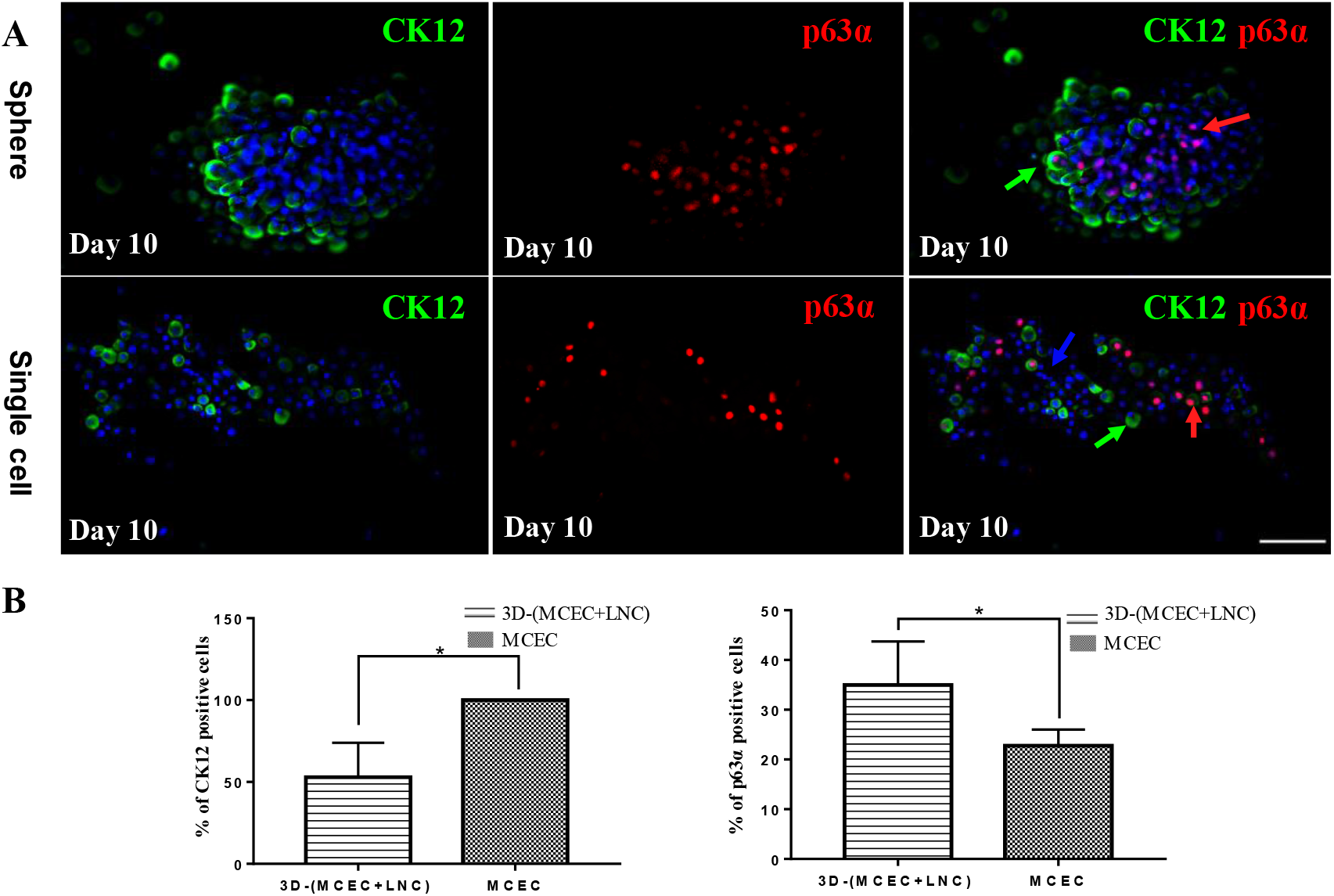
The expression changes of CK12 and p63α in 3D-MCEC+LNC. All spheres in 3D-MCEC+LNC group were collected on day10 It showed that more epithelial progenitor cell emerged in the center of the sphere and mature epithelial cells were in the outer part of the sphere. (Fig.5A, red arrow points to epithelial progenitor cell (p63α+), green arrow points to mature epithelial cells (CK12+), scale bar=100μm). Then spheres were digested into single cells by T/E method, the expression of CK12 and p63α in 3D-MCEC+LNC group was 52.95±20.99% and 35.58±8.11% respectively. (Fig.5A, lower row) Comparing with the MCEC group, the 3D-MCEC+LNC group showed increase of p63α expression, and the difference was statistically significant (P=0.043), among which the proportion of p63α+/CK12-cells were 11.51±3.76%. Comparing with the MCEC group, the positive rate of CK12 in the 3D-MCEC+LNC group reduced (P=0.000). (Fig.5B)

**Figure 6.**
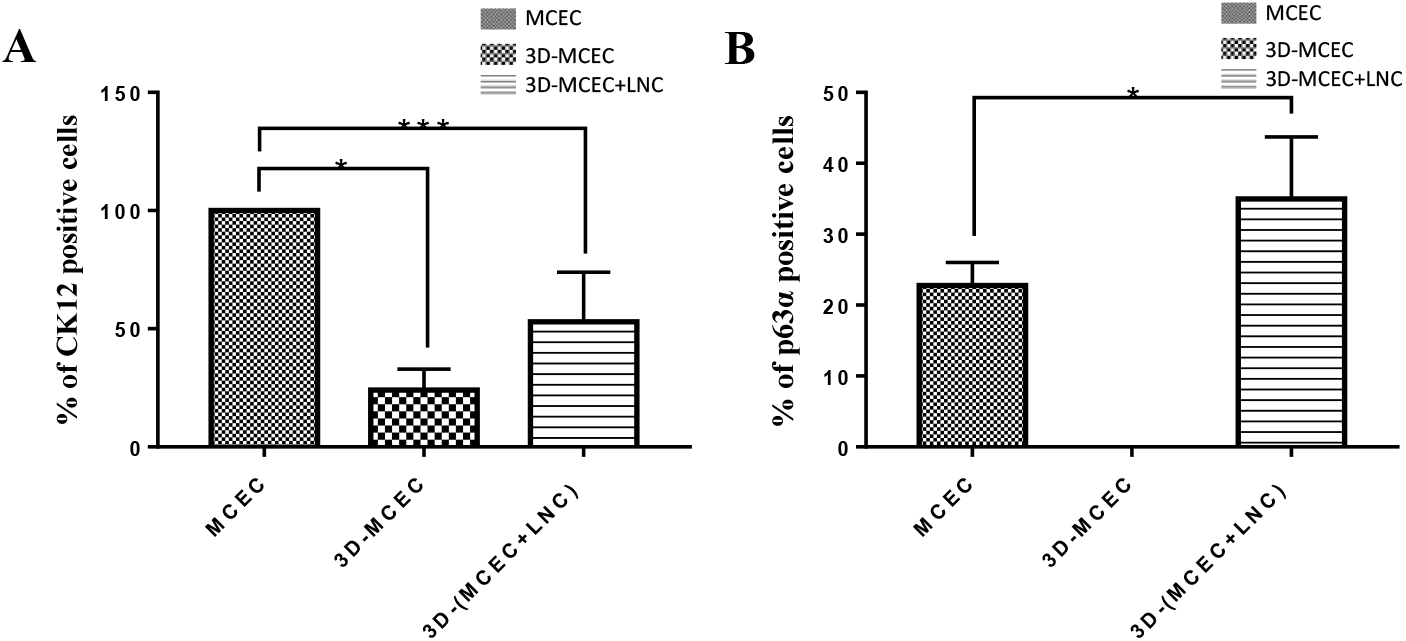
The expression changes of CK12 and p63α in three groups. There was no statistically significant difference in CK12 positive rate between the 3D-MCEC+LNC group and the 3D-MCEC group (P=0.051). (Fig.6A) Even though there was no p63α expression in 3D-MCEC group, comparing with the MCEC group, the 3D-MCEC+LNC group showed increase of p63α expression, and the difference was statistically significant (P=0.043). (Fig.6B)

### Spheres in 3D-MCEC+LNC group were larger than that in 3D-MCEC group

Measuring by ImageJ, it showed that the maximum diameter of the sphere was 180.32μm, the minimum was 52.01μm, and the average diameter was 109.39± 34.85μm in 3D-MCEC group. Whereas, In 3D-MCEC+LNC group, the maximum diameter was 306.20μm, the minimum diameter was 129.06μm and the average diameter was 212.24±57.91μm. (Fig.7 Table A) The diameter difference of cell sphere between the 3D-MCEC group and the 3D-MCEC+LNC group was statistically significant (P=0.000). (Fig.6 B)

**Figure 7.**
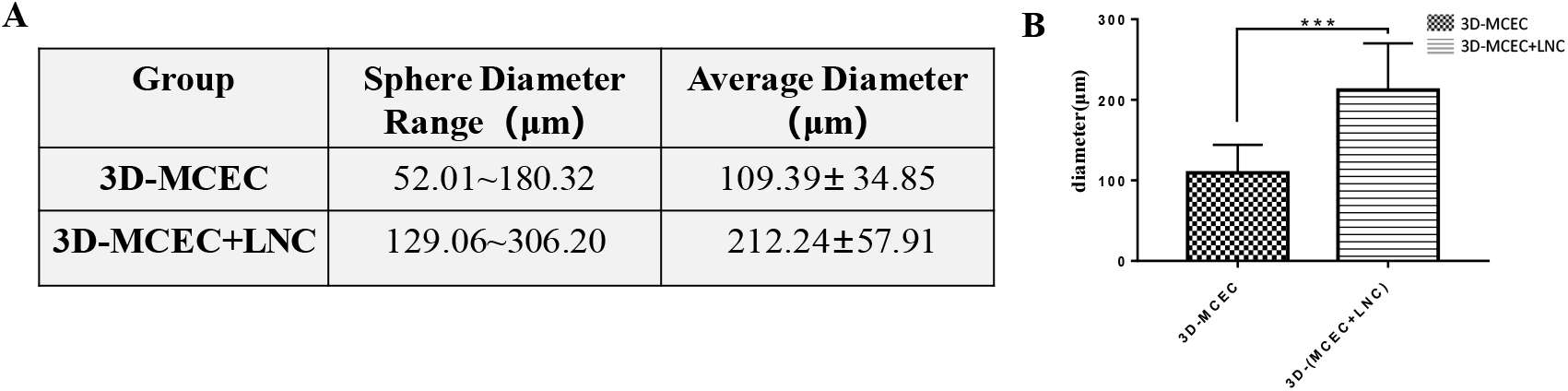
Sphere diameter of 3D-MCEC and 3D-MCEC +LNC group on day10 and expression of CK12/ p63α among three groups. In 3D-MCEC group, the minimum was 52.01μm, the maxim diameter of sphere was 180.32μm, and the average diameter was 109.39± 34.85μm. In 3D-MCEC+LNC group, the minimum diameter was 129.06μm, the maximum diameter was 306.20μm, and the average diameter was 212.24±57.91μm. (Fig.7 Table A) The diameter of cell sphere was compared between the 3D-MCEC +LNC group and the 3D-MCEC group, and the difference was statistically significant (P=0.000). (Fig.7B)

## DISCUSSION

Transcription factor p63 plays an important role in the proliferation and differentiation of epithelial cells, including three subtypes: α, β, γ, among which p63α is relatively expressed as the marker of LSC^21, 22^. Cytokeratin 12 (CK12) is considered as mature cornea epithelium cell specific marker^23^. In this study, we also observed that epithelial cells in the basal part of MCEC expressed p63α, but its positive rate was lower than that in limbal epithelium, which was consistent with the reports by Dua^24^ and Chang^25^. Our study demonstrated that p63α is not merely expressed in the basal part of the corneal limbus, but also in the basal layer of the central corneal epithelium. Thoft^26^ has proposed the “x-y-z” hypothesis to describe the maintenance of corneal epithelium in which X (proliferation of corneal epithelial basal cells) +Y (centripetal movement of cells) =Z(cell loss from the surface).The LSC located at the limbus of the cornea first differentiated into Transient Amplifying Cells (TAC), which migrated to the central cornea at the basal layer, and then differentiated into MCEC, supplementing the corneal epithelium^15^, suggesting that p63α may be a common marker of LSC and TAC, but the central corneal epithelium still has different levels of TAC, which is less mature and can express p63α. Majo^27^ and Vanclair^28^ also suggested that corneal epithelial stem cells may be found in the entire corneal epithelium of mammals, but this conclusion remains controversial.

3D Matrigel culturing is one of the best ways to simulate the microenvironment of stem cells in vivo, because it can make it more closed to the real state for cell morphology, proliferation, differentiation and material transportation ^29, 30^. In this study, MCECs were cultured in 3D Matrigel alone. After 10 days, the expression rate of CK12 in MCEC was significantly reduced, but the original expression of p63α in the basal layer part also disappeared. Whether CK12-/p63α-cells were corneal epithelial cells or transformed into other cells needs further study, but these results indicated that 3D Matrigel environment could reduce the maturity of MCEC, but not necessarily reverse differentiation into LSC.

The ability to form spheres growth in 3D Matrigel is regarded as a characteristic of neural stem cells^31^. Toshiro Sato^19^ cultured immortalized stem cell lines and primary cell lines isolated from tissues in 3D Matrigel, and then added into various growth factors and small molecular compounds to induce differentiation of organoid. Purshothama^13^ labeled the secretory cells of trachea epithelium of mice with yellow fluorescent protein (YFP). *In vitro* experiments, YFP+ secretory cells were cultured in the 3D Matrigel alone, with the YFP+ secretory cells mixed with epithelial basal cells as control group. Immunofluorescence staining showed that the YFP+ secretory epithelial cells could not only form cell spheres, but also expressed markers CK5 and p63α of epithelial basal stem cells, indicating that these mature secretory epithelial cells had reverse differentiation in vitro. However, CK5 and p63α were not expressed in YFP+ secreted epithelial cells in the control group, which was speculated by the authors that the presence of epithelial basal stem cells inhibited the reverse differentiation of mature cells. Jinny Jung Yoon^32^ conducted 3D sphere growth experiments using limbal cells digested by collagenase and proved that cells could aggregate and proliferate to form spheres. They used EdU and Ki-67 double-staining to show that the cells in the center of the sphere remained relatively inactive, but cell division became vigorous after they migrating out of the sphere. In addition, some cells expressed neuroprogenitor cell markers (nestin), myofibroblast markers (SMα), and limbal stem cell markers (p63α).The expression of Vimentin, K3 and K76 in the cell sphere could be stimulated by the culture of pro-differentiation medium, suggesting that the cell sphere was composed of a group of cells with different lineage and different degrees of differentiation.

In this study, we mixed LNC and MCEC in 3D Matrigel, the results showed that epithelial cells in spheres expressed less CK12 and more p63α comparing with MCEC and the ratio of p63α+/CK12-cell increased. Previous studies confirmed that LNC could maintain the undifferentiated state of LEPC^16, 17, 20^. Tseng SC^33, 34^ used the rabbit model to prove that when corneal epithelial tissue was combined with living limbal Matrigel, corneal epithelial cells lose mature phenotype (CK3), suggesting that limbal Matrigel microenvironment could induce the reverse differentiation of corneal epithelial cells to LSC. In this study, for the first time we demonstrated that LNC could decrease the expression of CK12 and increase the expression of p63α in MCEC, which indicated a tendency for MCEC to dedifferentiate into stem cells in 3D Matrigel environment with the help of LNC. Numerous other studies^35^ have reported the transdifferentiation of corneal epithelial cells, which have the potential to transform into fibroblasts in the Matrigel environment. Katikireddy^36^ et al. also found that both corneal stromal cells and BMSCs could be induced into limbal epithelial progenitor cells in a 3D environment, and differentiation efficiency of corneal stromal cells was higher. These studies have shown that corneal epithelial cells could transform phenotypes and antigen expression according to different microenvironments.

In this study, data shows that time needed for sphere growth formation in 3D-MCEC group was longer than that in 3D-MCEC+ LNC, and the sphere formed in 3D-MCEC+ LNC was larger and rounder. Previous studies have confirmed that PCK+ epithelial cells obtained from cornea limbus isolated by Dispase II and LNC (Vim+) could gather to form spheres in 3D Matrigel environment ^16, 17, 20, 37^. Xie^38^ confirmed that this cell aggregation was regulated by SDF-1/CXCR4 signaling, because SDF-1 was expressed in limbal epithelial cells, and CXCR4 was expressed in LNC. After the addition of CXCR4 inhibitors (AMD3100) in the culture environment, the cells of PCK+ and Vim+ no longer converged, and the resulting spheres were more differentiated and did not produce Holoclone on the 3T3 layer feeder. However, SDF-1 antigen was expressed in limbal epithelium and limbal stromal cells, but not in mature corneal epithelium and corneal stroma. The limbal stromal cells showed high expression of CXCR4 and weak expression in other parts. Therefore, we speculated that there might be other chemokines and related signaling pathway regulation for LNC to attract MCEC. Hemopoietic stem cell markers (CD117) and its ligand (SCF) are essential to regulating cell proliferation, differentiation and migration, and we reported^18^ previously that LNC could express SCF more than BMMSC, and limbal epithelial cells showed more differentiation antigen phenotype in mixture with LNC after the knock-out of SCF or blockade of the SCF/CD117 signaling pathways in 3D Matrigel. Therefore, SCF/CD117 signaling regulation may play a significant part in the co-culture of LNC and MCEC. Besides, we recently reported that the different gene expression between LNC and BMMSC^39^, which also suggested that LNC exhibit differential gene expression from BMMSC in the extracellular Matrigel (ECM) receptor interaction pathway and Wnt signaling pathway, indicating that LNC have their unique signaling pathways to support limbal stem cell niches. However, more studies need to further investigate the mechanisms of cell phenotype changes in 3D environment and it should be recognized that the process of dedifferentiation is also an inevitable part of tissue transformation and disease occurrence.

In conclusion, our study revealed for the first time that mature cornea epithelial cells obtained from central cornea lost mature cornea epithelial marker and gained more epithelial stem cell marker after ten days culture in 3D Matrigel with LNC. Therefore, mature cornea epithelial cells have the potential of dedifferentiation to cornea epithelial stem cell in the 3D Matrigel culture environment with the help of LNC, which could be a novel source of LSC for an alternative stem cell therapy for the treatment of LSCD. Further investigations are warranted to provide a more comprehensive understanding of potential mechanisms for the mature cornea epithelial cells dedifferentiation phenomenon.

## Supporting information

Supplementary Table S1. Materials Used for Tissue Isolation and Cell Culturing

## ACKNOWLEDGMENT

The authors wish to thank Dr Scheffer C. G. Tseng for the preparation of this manuscript.

## REFERENCES

1. Flaxman SR, Bourne RRA, Resnikoff S, et al. Global causes of blindness and distance vision impairment 1990-2020: a systematic review and meta-analysis. The Lancet Global Health 2017;5:e1221–e1234.

2. Xie LX, Shi WY The status quo and expectation of corneal research in China. Chinese Journal of Ophthalmology 2014;050:641–645.

3. Davanger M EA. Role of the pericorneal papillary structure in renewal of corneal epithelium. Nature 1971;229:560–561.

4. Nowell CS, Radtke F. Corneal epithelial stem cells and their niche at a glance. J Cell Sci 2017;130:1021–1025.

5. Chen JJ TS. Corneal epithelial wound healing in partial limbal deficiency. Invest Ophthalmol Vis Sci 1990;31:1301–1304.

6. Tseng SC. Concept and application of limbal stem cells. Eye(Lond) 1989;3(Pt 2):141–157.

7. Rama P, Matuska S, Paganoni G, Spinelli A, De Luca M, Pellegrini G. Limbal stem-cell therapy and long-term corneal regeneration. N Engl J Med 2010;363:147–155.

8. Vazirani J, Mariappan I, Ramamurthy S, Fatima S, Basu S, Sangwan VS. Surgical Management of Bilateral Limbal Stem Cell Deficiency. Ocul Surf 2016;14:350–364.

9. Kolli S, Ahmad S, Mudhar HS, Meeny A, Lako M, Figueiredo FC. Successful application of ex vivo expanded human autologous oral mucosal epithelium for the treatment of total bilateral limbal stem cell deficiency. Stem Cells 2014;32:2135–2146.

10. Utheim TP. Concise review: transplantation of cultured oral mucosal epithelial cells for treating limbal stem cell deficiency-current status and future perspectives. Stem Cells 2015;33:1685–1695.

11. Lin B, Srikanth P, Castle AC, et al. Modulating Cell Fate as a Therapeutic Strategy. Cell Stem Cell 2018.

12. Del Rio-Tsonis K, Tsonis PA. Eye regeneration at the molecular age. Dev Dyn 2003;226:211–224.

13. Tata PR, Mou HM, Pardo-Saganta A, et al. Dedifferentiation of committed epithelial cells into stem cells in vivo. Nature 2013;503:218–+.

14. Nasser W, Amitai-Lange A, Soteriou D, et al. Corneal-Committed Cells Restore the Stem Cell Pool and Tissue Boundary following Injury. Cell reports 2018;22:323–331.

15. Li W, Hayashida Y, Chen YT, Tseng SC. Niche regulation of corneal epithelial stem cells at the limbus. Cell Res 2007;17:26–36.

16. Li GG, Chen SY, Xie HT, Zhu YT, Tseng SC. Angiogenesis potential of human limbal stromal niche cells. Investigative ophthalmology & visual science 2012;53:3357–3367.

17. Li GG, Zhu YT, Xie HT, Chen SY, Tseng SC. Mesenchymal stem cells derived from human limbal niche cells. Invest Ophthalmol Vis Sci 2012;53:5686–5697.

18. Li G, Zhang Y, Cai S, et al. Human limbal niche cells are a powerful regenerative source for the prevention of limbal stem cell deficiency in a rabbit model. Scientific reports 2018;8:6566.

19. Sato T, Clevers H. SnapShot: Growing Organoids from Stem Cells. Cell 2015;161:1700–1700 e1701.

20. Xie HT, Chen SY, Li GG, Tseng SC. Isolation and expansion of human limbal stromal niche cells. Invest Ophthalmol Vis Sci 2012;53:279–286.

21. Pellegrini G, Dellambra E, Golisano O, et al. p63 identifies keratinocyte stem cells. Proceedings of the National Academy of Sciences of the United States of America 2001;98:3156–3161.

22. Di Iorio E, Barbaro V, Ruzza A, Ponzin D, Pellegrini G, De Luca M. Isoforms of DeltaNp63 and the migration of ocular limbal cells in human corneal regeneration. Proceedings of the National Academy of Sciences of the United States of America 2005;102:9523–9528.

23. Liu CY, Zhu G, Converse R, et al. Characterization and chromosomal localization of the cornea-specific murine keratin gene Krt1.12. The Journal of biological chemistry 1994;269:24627–24636.

24. Dua HS, Miri A, Alomar T, Yeung AM, Said DG. The role of limbal stem cells in corneal epithelial maintenance: testing the dogma. Ophthalmology 2009;116:856–863.

25. Chang CY, McGhee JJ, Green CR, Sherwin T. Comparison of stem cell properties in cell populations isolated from human central and limbal corneal epithelium. Cornea 2011;30:1155–1162.

26. Thoft RA, Friend J. The X, Y, Z hypothesis of corneal epithelial maintenance. Investigative ophthalmology & visual science 1983;24:1442–1443.

27. Majo F, Rochat A, Nicolas M, Jaoude GA, Barrandon Y Oligopotent stem cells are distributed throughout the mammalian ocular surface. Nature 2008;456:250–254.

28. Vauclair S, Majo F, Durham AD, Ghyselinck NB, Barrandon Y, Radtke F. Corneal epithelial cell fate is maintained during repair by Notch1 signaling via the regulation of vitamin A metabolism. Developmental cell 2007;13:242–253.

29. Brizzi MF, Tarone G, Defilippi P. Extracellular Matrigel, integrins, and growth factors as tailors of the stem cell niche. Curr Opin Cell Biol 2012;24:645–651.

30. Pera MF, Tam PP. Extrinsic regulation of pluripotent stem cells. Nature 2010;465:713–720.

31. Campos LS. Neurospheres: insights into neural stem cell biology. Journal of neuroscience research 2004;78:761–769.

32. Yoon JJ, Wang EF, Ismail S, McGhee JJ, Sherwin T. Sphere-forming cells from peripheral cornea demonstrate polarity and directed cell migration. Cell biology international 2013;37:949–960.

33. Kawakita T, Espana EM, He H, Li W, Liu CY, Tseng SC. Intrastromal invasion by limbal epithelial cells is mediated by epithelial-mesenchymal transition activated by air exposure. Am J Pathol 2005;167:381–393.

34. Espana EM, Kawakita T, Romano A, et al. Stromal niche controls the plasticity of limbal and corneal epithelial differentiation in a rabbit model of recombined tissue. Invest Ophthalmol Vis Sci 2003;44:5130–5135.

35. Kalluri R, Neilson EG. Epithelial-mesenchymal transition and its implications for fibrosis. J Clin Invest 2003;112:1776–1784.

36. Katikireddy KR, Dana R, Jurkunas UV. Differentiation potential of limbal fibroblasts and bone marrow mesenchymal stem cells to corneal epithelial cells. Stem cells 2014;32:717–729.

37. Han B, Chen SY, Zhu YT, Tseng SC. Integration of BMP/Wnt signaling to control clonal growth of limbal epithelial progenitor cells by niche cells. Stem cell research 2014;12:562–573.

38. Xie HT, Chen SY, Li GG, Tseng SC. Limbal epithelial stem/progenitor cells attract stromal niche cells by SDF-1/CXCR4 signaling to prevent differentiation. Stem cells 2011;29:1874–1885.

39. Wang W, Li S, Xu L, et al. Differential Gene Expression between Limbal Niche Progenitors and Bone Marrow Derived Mesenchymal Stem Cells. Int J Med Sci 2020;17:549–557.

