## Supplementary Table S1. Materials Used for Tissue Isolation and Cell Culturing for "Limbal Niche Cells in Three-Dimensional Matrigel Induced Dedifferentiation of Mature Corneal Epithelial Cells toward a Progenitor State"

**SUPPLEMENTARY** MATERIAL

| **Supplementary Table S1. Materials Used for Tissue Isolation and Cell Culturing** | | |
| --- | --- | --- |
| **Materials** | **Sources** | **Concentration** |
| Dulbecco's modified Eagle's Medium (DMEM) | Invitrogen, Grand Island, NY | DMEM/F-12 (1:1) |
| F-12 nutrient mixture (F-12) | Invitrogen, Grand Island, NY | DMEM/F-12 (1:1) |
| Knockout Serum Replacement | Invitrogen, Grand Island, NY | 10% |
| Fetal Bovine Serum (FBS) | Invitrogen, Grand Island, NY | 10% |
| Human Fibroblast Growth Factor-Basic | Invitrogen, Grand Island, NY | 4 ng/ml |
| Leukemia inhibitory factor (LIF) | Invitrogen, Grand Island, NY | 10 ng/ml |
| Dimethyl Sulfoxide | Sigma-Aldrich, St. Louis, MO | 0.5% |
| Insulin-Transferrin-sodium selenite media supplement | Roche, Indianapolis, IN | 5 µg/ml insulin, 5 µg/ml Transferrin, 5 ng/ml sodium selenite |
| Hank's Balanced Salt Solutions (HBSS) | Invitrogen, Grand Island, NY | 1 x |
| Phosphate-Buffered Saline pH7.4 (PBS) | Invitrogen, Grand Island, NY | 1 x |
| Amphotericin B | Invitrogen, Grand Island, NY | 50 µg/ml |
| Gentamicin | Invitrogen, Grand Island, NY | 1.25 µg/ml |
| Dispase II | Roche, Indianapolis, IN | 10 mg/ml |
| Collagenase A | Roche, Indianapolis, IN | 1 mg/ml |
| Cultrex 3-D Culture Matrix™ Basement Membrane Matrix | Trevigen, Gaithersburg, MD | 50% |
| Trypsin and EDTA (T/E) | Invitrogen, Grand Island, NY | 0.25% and 1 mM |

| **Supplementary Table S2: Antibodies Used for Immunofluorescence Staining (IF)** | | |
| --- | --- | --- |
| **Antibodies** | **Sources** | **Dilution** |
| Cytokeratin 12 (L-20) | Santa Cruz (Santa Cruz, CA) | 1:50 |
| p63α | Cell Signaling (Danvers, MA) | 1:400 |
| Pan-cytokeratins (AE1/AE3) (PCK) | Abcam (Cambridge, MA) | 1:200 |
| Vimentin (SP20) | Abcam (Cambridge, MA) | 1:400 |
| Alexa Fluor 488 anti–donkey IgG | Invitrogen (Carlsbad, CA) | 1:200 |
| Alexa Fluor 594 anti–donkey IgG | Invitrogen (Carlsbad, CA) | 1:200 |
| DAPI | Sigma-Aldrich (St. Louis, MO) | 1:100 |
